# Targeting processive transcription for Myc-driven circuitry in medulloblastoma

**DOI:** 10.1101/2025.03.14.643337

**Authors:** Lays Martin Sobral, Faye M. Walker, Krishna Madhavan, Elizabeth Janko, Sahiti Donthula, Ilango Balakrishnan, Dong Wang, Angela Pierce, Mary M. Haag, Billie J. Carstens, Natalie J. Serkova, Nicholas K. Foreman, Sujatha Venkataraman, Bethany Veo, Rajeev Vibhakar, Nathan A. Dahl

**Author notes:** These authors contributed equally to this work (Lays Martin Sobral, Faye M. Walker, Krishna Madhavan, Rajeev Vibhakar, Nathan A. Dahl). Corresponding Author: Nathan A. Dahl, MD, Department of Pediatrics, University of Colorado School of Medicine, Mail Stop # 8302 Research Complex 1, North Tower, 12800 E. 19th Ave., Rm P18-4118, Aurora, CO 80045. |.

## Abstract

**Background:** Medulloblastoma is the most common malignant brain tumor of childhood. The highest-risk tumors are driven by recurrent Myc amplifications (Myc-MB) and experience poorer outcomes despite intensive multimodal therapy. The Myc transcription factor defines core regulatory circuitry for these tumors and acts to broadly amplify downstream pro-survival transcriptional programs. Therapeutic targeting of Myc directly has proven elusive, but inhibiting transcriptional cofactors may present an indirect means of drugging the oncogenic transcriptional circuitry sustaining Myc-MB.

**Methods:** Independent CRISPR-Cas9 screens were pooled to identify conserved dependencies in Myc-MB. We performed chromatin conformation capture (Hi-C) from primary patient Myc-MB samples to map enhancer-promoter interactions. We then treated *in vitro* and xenograft models with CDK9/7 inhibitors to evaluate effect on Myc-driven programs and tumor growth.

**Results:** Eight CRISPR-Cas9 screens performed across three independent labs identify CDK9 as a conserved dependency in Myc-MB. Myc-MB cells are susceptible to CDK9 inhibition, which is synergistic with concurrent inhibition of CDK7. Inhibition of transcriptional CDKs disrupts enhancer-promoter activity in Myc-MB and downregulates Myc-driven transcriptional programs, exerting potent anti-tumor effect.

**Conclusions:** Our findings identify CDK9 inhibition as a translationally promising strategy for the treatment of Myc-MB.

**Key Points:**

- CDK9 is an intrinsic dependency in Myc-driven medulloblastoma
- Dual CDK9/7 inhibition disrupts Myc-driven transcriptional circuitry
- CDK9 inhibitors should be developed as pharmaceutical agents for Myc-MB

**Importance of the Study:** Medulloblastoma is the most common malignant brain tumor of childhood, and outcomes for high-risk subgroups remain unsatisfactory despite intensive multimodal therapy. In this study, we pool multiple independent CRISPR-Cas9 screens to identify transcriptional cofactors such as CDK9 as conserved dependencies in Myc-MB. Using Hi-C from primary patient samples, we map Myc enhancer-promoter interactions and show that they can be disrupted using inhibition of transcriptional CDKs. CDK9 inhibitor treatment depletes Myc-driven transcriptional programs, leading to potent anti-tumor effect *in vitro* and prolongation of xenograft survival *in vivo*. With a large number of CDK9 inhibitory compounds now in clinical development, this study highlights the opportunity for clinical translation of these for children diagnosed with Myc-MB.

## Introduction

Medulloblastoma (MB) is the most common malignant brain tumor in children^1^. Great progress has been made in defining molecular subclassifications^2–4^, understanding developmental origins^5,6^, and stratifying multimodal therapy^7,8^ for this heterogenous family of tumors, and the majority of children diagnosed with medulloblastoma today are now cured of their disease^7,8^. However, high-risk subgroups continue to experience unsatisfactory outcomes, particularly for patients with group 3 tumors driven by *MYC* amplification (Myc-MB). Patients with Myc-MB experience higher rates of metastasis, increased likelihood of relapse, shorter time to death following relapse, and poorer rates of overall survival^9,10^. While intensification of conventional cytotoxic chemotherapy has improved outcomes for these patients^7^, no clinical strategies to directly or indirectly target Myc as the inciting oncogenic driver currently exist.

Myc is a transcription factor essential for the regulation of proliferative potential in normal progenitor tissues, and it is co-opted as a driver through amplification or other overexpression across a range of cancer types^11–13^. Transcription of the *MYC* locus itself is regulated through a combinatorial array of distal enhancer elements, non-coding transcriptionally active loci that regulate and amplify transcription at a target promoter^12,14^. Deletion of these enhancer clusters, and thus loss of the feed-forward transcriptional amplification, leads to a complete loss of Myc expression comparable to Cre-mediated conditional deletion of the *MYC* locus itself^12^. In parallel, disruption of transcriptional machinery preferentially effects genes with strong association to super-enhancer (SE) elements, such as *MYC*^15^. This has led numerous groups to investigate targeting various components of transcriptional machinery as a means of drugging Myc indirectly^13,15,16^.

Downstream, the Myc transcription factor acts to broadly amplify transcriptional activity, both at established Myc target genes and at pre-existing expression programs defined by cell identity and state^11^. Once localized to chromatin, Myc acts not to facilitate RNA polymerase II (Pol II) recruitment and initiation, but rather to drive release of paused Pol II and productive elongation into gene bodies^17^. Thus, the diverse array of functional programs which fall under Myc regulation are, to a significant degree, contingent on its ability to amplify transcriptional kinetics at the step of elongation checkpoint control^17,18^.

Processive transcription by Pol II is controlled through dynamic phosphorylation of the highly conserved C-terminal domain (CTD)^19–21^. Following Pol II recruitment, initiation is governed by the pre-initiation complex, in which the CDK7-containing TFIIH phosphorylates the CTD at the Ser5 position^22^. After transcribing 20-80 bases downstream, Pol II then becomes paused in promoter-proximal regions at many genomic loci. Phosphorylation of the CTD Ser2 by the CDK9/CyclinT1 pair, collectively termed positive transcription elongation factor b (P-TEFb), is then required for pause release^21,23,24^. P-TEFb is carried from an inactive reservoir pool to chromatin by a variety of bromodomain and extraterminal (BET) family and super elongation complex (SEC) proteins, enabling specificity in targeting P-TEFb recruitment to genomic loci^25^. Degradation or inhibition of these P-TEFb containing complexes has been shown to downregulate Myc expression and Myc-driven transcriptional programs^13,16,26,27^. Much of this biology has been dissected, however, using chemical tool compounds^13,15,16,26^ without potential for implementation as clinical grade pharmaceuticals and with little data regarding penetration across the blood-brain-barrier.

In this study, we pool multiple independent CRISPR-Cas9 screens to identify transcriptional CDKs as conserved dependencies in Myc-MB. We examine their role in sustaining Myc-driven transcriptional circuitry, and we explore whether clinically relevant small molecule inhibitors could be used to leverage this dependency as a translationally actionable Myc-targeted therapy for medulloblastoma.

## Materials and Methods

### Western Blotting

Western blotting was performed as previously described^28^. Antibodies used were as follows, prepared according to manufacturer recommendations: CDK9 (C12F7) Rabbit mAb Cell Signaling #2316, c-Myc (D84C12) Rabbit mAb Cell Signaling #5605, β-Actin (13E5) Rabbit mAb Cell Signaling #4970, Phospho-Rpb1 CTD (Ser2) (E1Z3G) Rabbit mAb Cell Signaling #13499, Phospho-Rpb1 CTD (Ser5) (D9N5I) Rabbit mAb Cell Signaling #13523, Rpb1 CTD (4H8) Mouse mAb Cell Signaling #2629, α-Tubulin (DM1A) Mouse mAb Cell Signaling #3873, Cleaved Caspase-3 (Asp175) Antibody Cell Signaling #9661, Anti-rabbit IgG, HRP-linked Antibody Cell Signaling #7074.

### Click-iT nascent RNA and Immunofluorescence

Click-iT reaction for nascent RNA (Click-iT RNA HCS Assay, C10327, Invitrogen) was performed according to manufacturer instructions following ZTR treatment or DMSO for 24 hrs as previously described^29^. Cells were then incubated with Myc antibody (Cell Signalingl, 5605S) and secondary antibody (Goat anti-Rabbit IgG Alexa Fluor 568, Invitrogen A-11011) prior to staining with HCS NuclearMask Blue stain solution. Images obtained using a confocal microscope (Keyence).

### shRNA Lentiviral Production

Lentiviral packaging was performed as previously described^29^. Control and shCDK9 plasmids were obtained from the University of Colorado Functional Genomics Core Facility (CDK9 #1: TRCN0000000494, CDK9 #2: TRCN0000000497, CDK9 #3: TRCN0000000498).

### Drug Synergy

Cells were seeded 1,000 cells/well in 96-well ultra-low-attachment round-bottom tissue culture plates (Corning) and centrifuged to promote neurosphere formation. Neurospheres were then treated with indicated concentrations of AZD4573 and SY5102 and imaged for one week using an IncuCyte S3 Live Cell Analysis System. Neurosphere measurements were then normalized to the control wells and analyzed per Bliss and HSA synergy models using Combenefit software (v2.021)^30^.

### Xenograft Models

Athymic nude mice aged 4 to 8 weeks were anesthetized, immobilized, and prepared for intracranial injection as previously described^28^. To target the cerebellum, a 1.0 mm diameter burr was drilled in the cranium with a dental drill bit at 1.500 mm to the right and 2.000 mm posterior to lambda. A suspension of D458 cells was prepared by suspending a concentration of 20,000 cells/3 μl in serum free media per injection, and slowly injected 3.000 mm ventral to the surface of the skull using an UltraMicroPump III and a Micro4 Controller (World Precision Instruments) equipped with a Hamilton syringe and a 26-gauge needle. The burr hole was sealed, incision closed, and topical antibiotic applied as previously described. Post-surgical pain was controlled with SQ carprofen (5 mg/kg/daily) on the day of and for 2 days following the procedure. Flank models generated as previously described^29^. University of Colorado Institutional Animal Care and Use Committee (IACUC) approval was obtained and maintained throughout the conduct of the study.

### CUT&RUN

<u>C</u>leavage <u>U</u>nder <u>T</u>argets & <u>R</u>elease <u>U</u>sing <u>N</u>uclease assays were performed as previously described^29^ using H3K4me3 (EpiChypher 13-0041) and Myc (Cell Signaling, 13987S) antibody in accordance with the manufacturer’s protocol (CUTANA ChIC/CUT&RUN EpiCypher 14-1048). The NEBNext Ultra DNA Library Prep Kit for Illumina (NEB #E7645S) with Dual Index Primers (NEB #E7600) were used for library preparation prior to sequencing on the NovaSeq6000 platform.

### Chromosome Conformation Capture (Hi-C)

Hi-C reactions and library preparations were performed by Arima Genomics. FFPE samples were sectioned at 5-10 µm thickness, de-waxed, and rehydrated prior to sample preparation using the Arima-HiC+ for FFPE kit (#A311038). Sequencing libraries were prepared by shearing proximally ligated DNA followed by size-selection of DNA fragments using SPRI beads. Size-selected fragments were then enriched for fragments containing ligation junctions using streptavidin beads before library construction. Illumina-compatible sequencing libraries were prepared using the Arima Library Prep Module (#A303011) according to manufacturer instructions and sequenced on the NovaSeq6000 platform.

After sample demultiplexing, raw read pairs were mapped to the reference genome (hg38) and de-duplicated using HiCUP^31^. Processed alignments are then converted into Hi-C matrices using Juicer v1.6^32^ and visualized using Juicebox^33^. Chromatin loop calls were identified using HiCCUPS algorithm with KR normalization. TADs were identified using Arrowhead algorithm at multiple resolutions.

## Results

### Integrated CRISPR-Cas9 Screening Identifies CDK9 as a Conserved Dependency in Myc-MB

We have previously performed and reported CRISPR-Cas9 screening of 1,140 druggable targets across three established models of Myc-MB^34^. In order to strengthen the reproducibility of this approach and further enrich for conserved dependencies to prioritize for translational development, we intersected these data with published CRISPR-Cas9 screens performed specifically in Myc-MB by two additional independent labs (eight cell lines total)^35,36^ (**Figure 1A** and **Supplemental Table 1**). As the statistical cutoffs applied and the detail of raw data reported varied between publications, we included significant hits as originally defined by the source manuscript authors. This resulted in 35 genes consistently identified as significant dependencies across all three screens (**Figure 1B**). Conserved hits included genes with well-characterized roles in cell cycle regulation (*CDK1/6, CHEK1, WEE1, AURKB, PLK4, CDC7*), nucleotide synthesis (*DHODH, RRM1/2, TYMS*), and DNA damage response (*POLE, MCL1, RAD51, BCL2L1*) (**Supplemental Table 1**). Also notable were hits for the transcription-associated cyclin-dependent kinases CDK9 and CDK8, with the closely related CDK7 scoring in 7 of 8 cell lines^34–36^.

**Figure 1.**
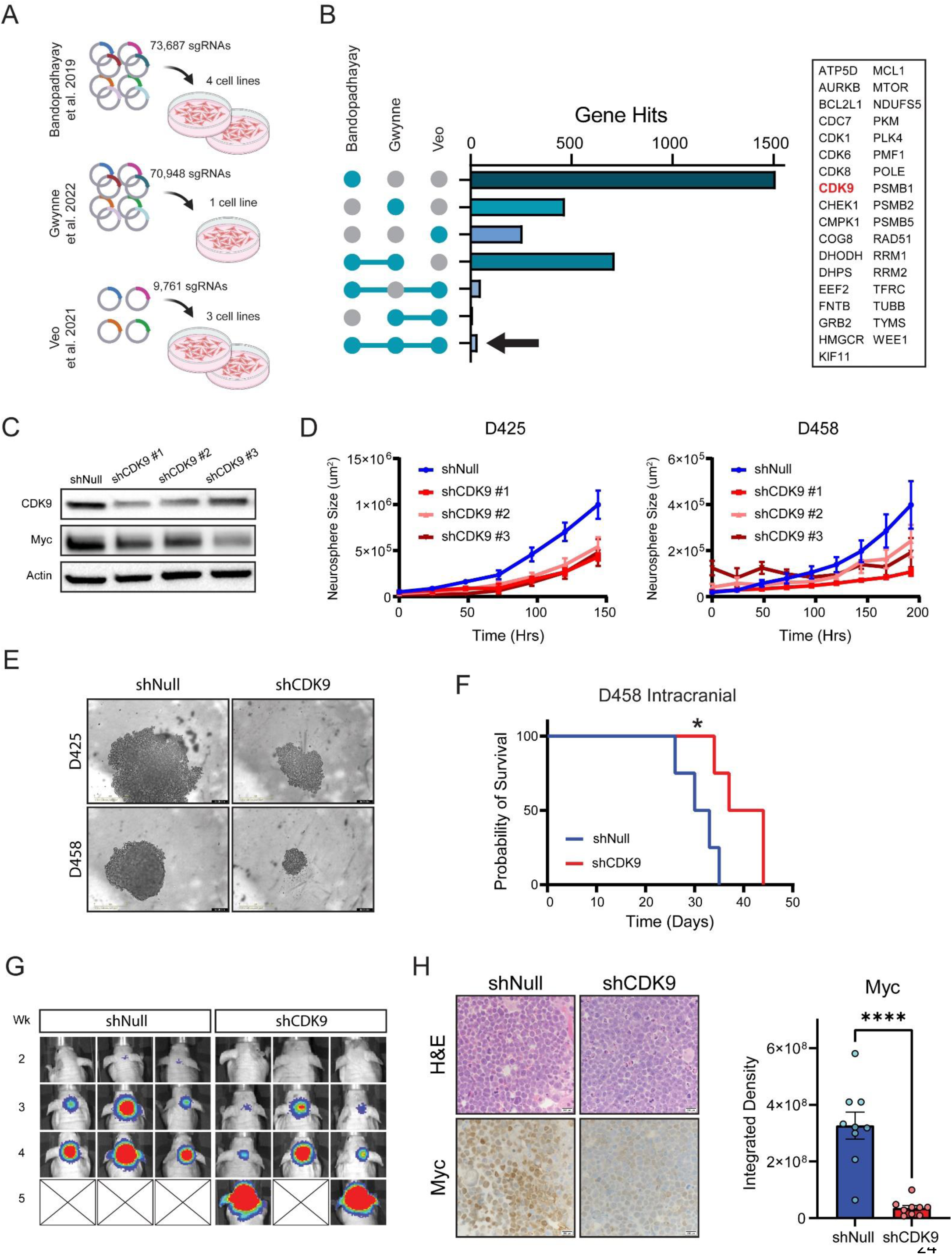
CDK9 is a conserved functional dependency in Myc-MB. **A.** Schematic representation of published CRISPR-Cas9 screens pooled for this analysis, with unique sgRNAs and cell lines indicated. **B.** UpSet plot (left) of unique and common hits from each independent screen. Arrow indicates genes scored as hits in all three screens, listed in table on right. **C.** Immunoblot in D458 cells following shNull or shCDK9 induction. **D.** Growth of D425 and D458 cells expressing shNull or shCDK9 in neurosphere conditions. **E.** Representative live cell images from (D). **F.** Kaplan-Meier survival analysis of D458 orthotopic xenografts expressing shNull or shCDK9 (log rank *p < 0.05). **G.** Representative bioluminescent imaging from animals in (F). **H.** Representative immunohistochemistry for Myc in tumors harvested from (f), quantification shown on right. Comparison reflects two-tailed Student’s t-test (**** p=<0.0001).

In order to validate CDK9 as a relevant therapeutic target in Myc-MB, we next transduced two established Myc-driven cell lines (D425 and D458) with non-overlapping shRNA constructs targeting CDK9 or scrambled control. Successful knockdown of CDK9 protein expression resulted in a proportional decrease in expression of Myc and a loss of proliferative potential when cells were cultured in neurosphere conditions (**Figure 1C-E**). We then engrafted the shNull-D458 or shCDK9-D458 cells orthotopically in the cerebellum of athymic nude mice. CDK9 knockdown led to a delay in tumor development and significant prolongation in animal survival (median survival 32 days shNull vs 41 days shCDK9, p=0.03) (**Figure 1F-G**). When tumors were collected at endpoint, shCDK9-D458 tumors showed considerably lower Myc expression by immunohistochemistry (**Figure 1H**). In total, these data identify CDK9 as a conserved dependency in Myc-driven medulloblastoma and suggest it may be amenable to targeting for anti-tumor effect.

### Myc-MB is Susceptible to Pharmacologic Inhibition of CDK9

There has been growing interest in CDK9 as a promising target for cancer therapy, leading to the formulation of increasingly specific inhibitory compounds now in preclinical and early clinical development^37,38^. We therefor sought to assess whether the above anti-tumor effects could be replicated using small molecule inhibitors targeting CDK9 (CDK9i). We tested a range of available CDK9i against a panel of Myc-driven (D425, D458) and non-Myc-driven (ONS76, UW228) medulloblastoma lines. We broadly observed nanomolar (atuveciclib, AZD4573, flavopiridol, PHA767491) or low-micromolar (LDC00067) susceptibility to CDK9i across the models tested, with a trend towards lower mean inhibitory concentrations in the Myc-MB (**Figure 2A**). We selected two compounds highly specific for CDK9 (vs other CDKs or kinases), atuveciclib^39^ and AZD4573^40^, for further testing (**Figure 2B**). Treatment of Myc-MB lines with either compound led to a marked reduction in Myc mRNA and protein expression (**Figure 2C-D**). When tested in neurosphere culture, a dose-dependent reduction in fitness was observed (**Figure 2E**), supporting the efficacy of CDK9-selective agents as anti-tumor therapies for Myc-MB.

**Figure 2.**
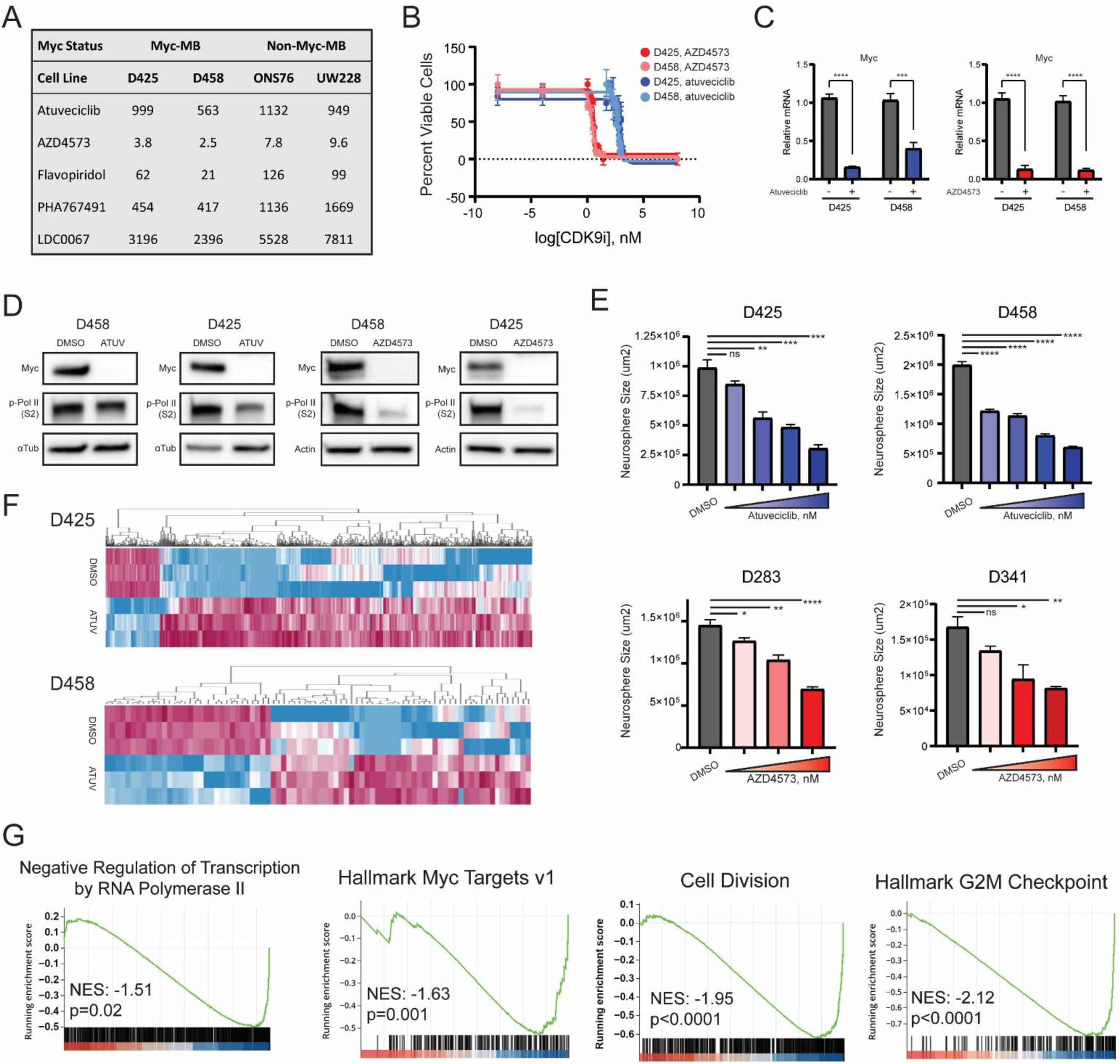
Myc-MB cells are susceptible to CDK9 pharmacologic inhibition. **A.** Screen of five commercially available CDK9i against Myc-(D425, D458) and non-Myc (ONS76, UW228) MB lines. **B.** Mean inhibitory concentrations of atuveciclib and AZD4573 in D425 and D458 cells. **C.** qPCR for Myc mRNA following atuveciclib or AZD4573 treatment. Comparison reflects two-tailed Student’s t-test (*** p=<0.001, **** p=<0.0001). **D.** Immunoblot for Myc and Ser2-phosphorylated Pol II following atuveciclib or AZD4573 treatment. **E.** Neurosphere size of indicated cells treated at increasing concentrations of atuveciclib (0 to 1000 nM) or AZD4573 (0 to 20 nM). Comparison reflects two-tailed Student’s t-test (* p=<0.05, ** p=<0.01, *** p=<0.001, **** p=<0.0001). **F.** Unsupervised hierarchical clustering of RNA-seq from D425 and D458 cells following atuveciclib treatment (n=3 biological replicates, log2FC > 2, variance > 1, and p-value < 0.01). **G.** Gene set enrichment analysis from (F).

To better understand the mechanistic consequences of selective CDK9 inhibition in Myc-MB, we then performed RNA sequencing (RNA-seq) in atuveciclib-treated cells compared to untreated controls in both D425 and D458 cell lines. This identified 532 - 1728 differentially expressed genes with pval <0.01 and LFC >1.2 (**Figure 2F**). Atuveciclib treatment resulted in a significant downregulation of both transcription by RNA Pol II and hallmark Myc target programs, as well as gene sets related to cell division and cell cycle progression (**Figure 2G**). Functional programs demonstrated strong overlap between the two models (**Supplemental Figure 1**). Taken together, these support CDK9 inhibition as an effective means of drugging transcriptional circuitry at the heart of Myc-driven programs.

### Dual Targeting of Processive Transcription Exhibits Therapeutic Synergy in Myc-MB

Regulation of processive transcription by RNA Pol II is not a single-step checkpoint but instead is governed by inputs from multiple transcriptional CDKs across the transcription cycle^19–21^. Our team has previously identified and characterized the role of the TFIIH-member CDK7 in Myc-driven circuitry within MB^34^. Given the closely related biological functions of these two kinases, we asked whether they could be targeted in parallel, both to augment anti-tumor effect and expand the repertoire of available small molecule inhibitors available for study. Again using AZD4573 as a CDK9-specific tool compound, we tested it at increasing concentrations with SY5102, a highly-selective noncovalent inhibitor of CDK7^41^. When assessing synergy using the Bliss model, we observed a remarkable synergistic effect at very low nanomolar concentrations (synergy defined as Bliss score ≥10, observed scores >30-40) (**Figure 3A**). Analysis using HSA modeling of synergy yielded similar positive scores (**Supplemental Figure 2**), supporting the utility of targeting these transcriptional cofactors in parallel.

**Figure 3.**
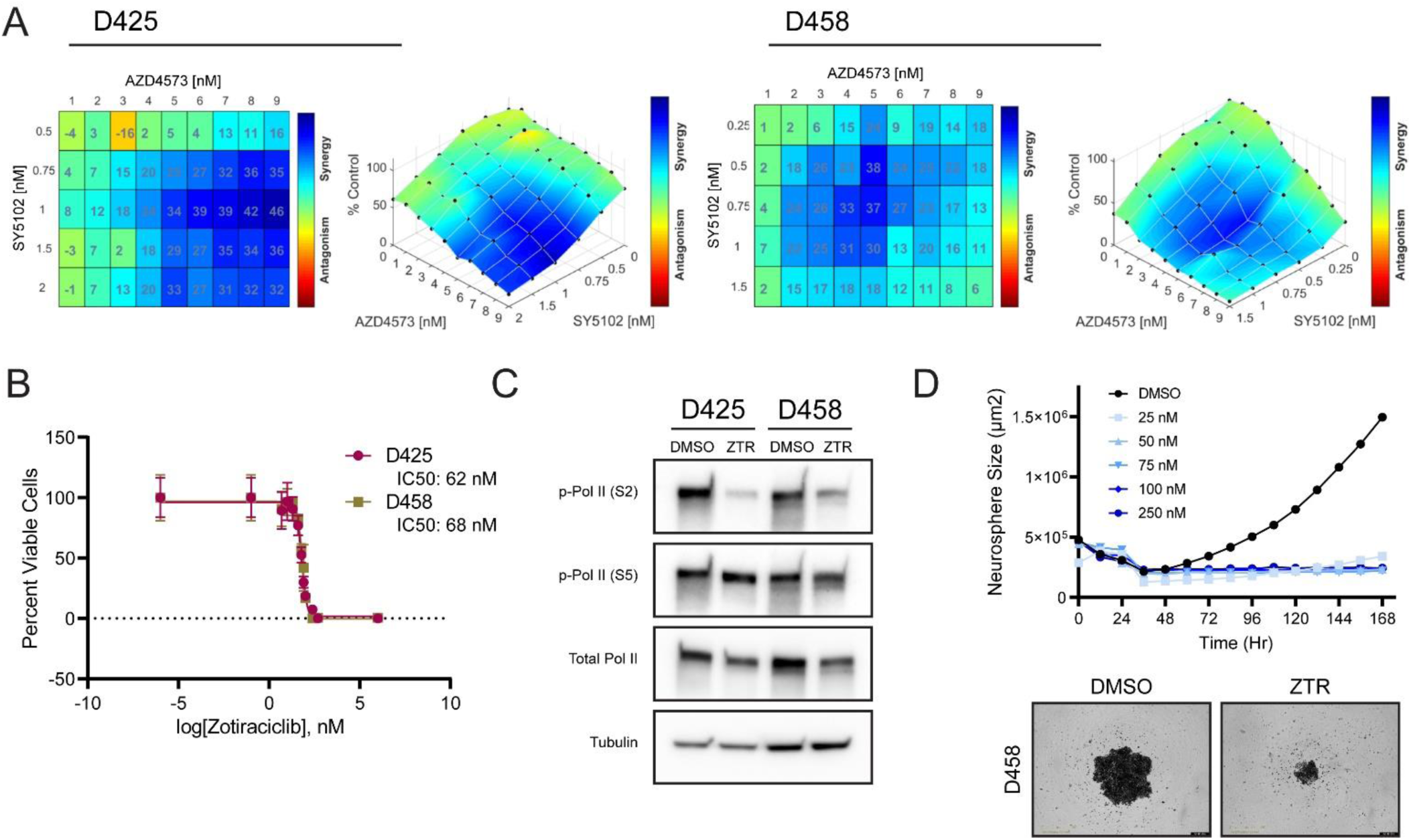
Combination CDK7i and CDK9i is synergistic in Myc-MB. **A.** Mapped Bliss synergy scoring at increasing concentrations of AZD4573 and SY5102 in D425 and D458 cells. Scores ≥10 are considered synergistic, represented as cyan to blue. **B.** Mean inhibitory concentrations of zotiraciclib in indicated Myc-MB models. **C.** Immunoblot of Ser2- and Ser5-phosphorylated Pol II following zotiraciclib treatment. **D.** D458 sphere size when grown in neurosphere conditions at indicated ZTR concentrations. Representative live cell imaging shown below.

Following this, we turned our attention to the drug zotiraciclib. Zotiraciclib (ZTR) is an orally-available, pyrimidine-based inhibitor that inhibits CDK9 at an IC50 of 3 nM and CDK7 at 37 nM (Cothera Bioscience, Investigator Brochure), and it is currently in phase I/II trials for adults with relapsed anaplastic astrocytoma or glioblastoma (^42^ and NCT03224104, NCT02942264). When tested against the D425 and D458 models of Myc-MB, it demonstrated anti-tumor effect with mean inhibitory concentrations in the low nanomolar range and inhibition of CDK-catalyzed Pol II CTD phosphorylation (**Figure 3B-C**). We then tested dose-dependent effects with Myc-MB cells cultured in neurosphere conditions. At concentrations tested as low as 25 nM, all ZTR dose exposure resulted in a complete cessation of growth in neurosphere media (**Figure 3D**). Together, this supported combined targeting of CDK9 with CDK7 as a promising means of disrupting transcriptional circuitry in Myc-MB.

### CDK9/7 Inhibition Ablates Enhancer-Promoter Activity within Myc-Amplified TAD

The *Myc* promoter is regulated by inputs from distinct, evolutionarily conserved super enhancers, and disruption of this enhancer activity leads to a loss of *Myc* expression comparable to deletion of the *Myc* locus itself^12,15,43^. Given the comparatively high levels of transcriptional output at enhancer or super enhancer loci, inhibition of transcriptional machinery has been shown to preferentially deplete enhancer-driven oncogenes^13,15^. Active enhancer loci or specific enhancer-promotor (E-P) regulatory looping are tightly regulated across development and cell fate commitment, but they can be altered in states of neoplastic transformation, including medulloblastoma^44–46^. This complicates experimental assessments of enhancer perturbation, as E-P inferences based on genomic distance or transcriptional correlation may be confounded by complex chromatin rearrangements.

To address this, we performed chromatin conformation capture (Hi-C) from two primary FFPE patient samples of Myc-MB^47^. Each patient sample was clinically annotated as a Group 3 tumor with Myc amplification in double minute pattern by FISH (**Figure 4A** and **Supplemental Figure 3**). Coverage maps delineated quite distinct regions of amplification between patients (approximately 5.7 Mb upstream in UPN1355, 885 Kb downstream in UPN1433), though both encompassed the entirety of the *Myc* gene body (**Figure 4B**). When examining loops anchored at the *Myc* promoter, we called 14 unique loops in one sample and 15 in the second. Of these, 8 were present in both patient samples, while the remainder were patient-specific (**Figure 4B** and **Supplemental Table 2**). This is consistent with prior reports that while promoters of core oncogenic drivers maintain similar active epigenetic signatures across a diagnostic entity, non-promotor regulatory inputs may vary widely from patient to patient^48^.

**Figure 4.**
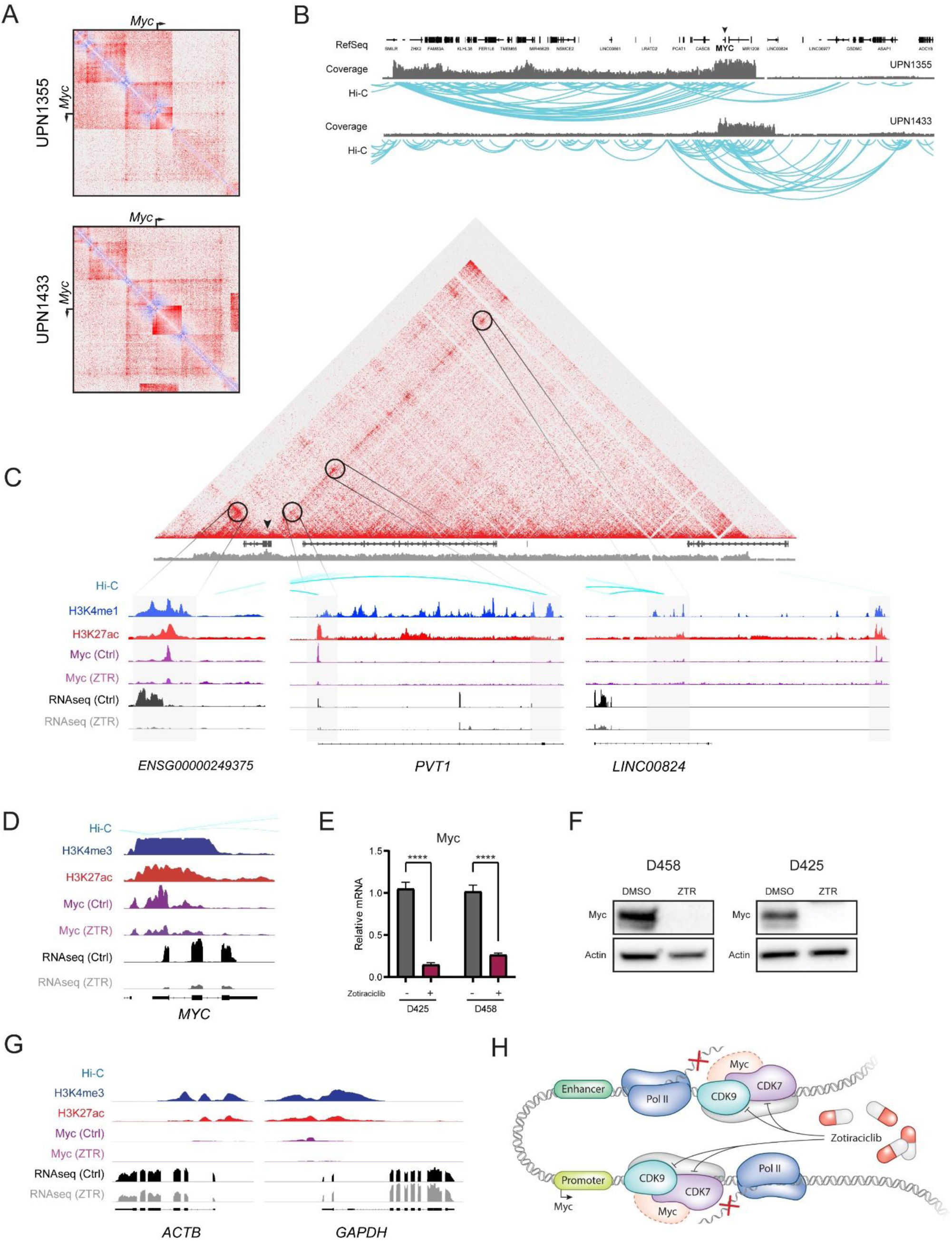
CDK9/7 inhibition disrupts *MYC* enhancer-promoter activity within the Myc-amplified TAD. **A.** Primary patient Myc-MB Hi-C contact maps centered at *MYC* promoter, heatmap visualized as observed over statistically expected contact frequency. **B.** Hi-C arc plot visualizing contact loops, again centered at *MYC* promoter (arrowhead). Contact loops demonstrate both conserved and patient-specific 3D interactions. Coverage map shows patient-specific region of amplification. **C.** UPN1433 Hi-C contact map centered on *MYC*-containing TAD, aligned with respective ChIP-seq or CUT&RUN tracks. Hi-C arcs below (cyan) denote contacts to *MYC* promoter, with other loops removed for clarity. Active enhancer elements are defined by H3K4me1, H3K27ac, and contact looping. Myc occupancy and RNA-seq transcript at each loci are shown before and after zotiraciclib treatment. **D.** Above data centered at annotated *MYC* promoter, marked by H3K4me3. Myc occupancy and RNA-seq transcript are shown before and after zotiraciclib treatment. qPCR (**E**) and immunoblot (**F**) for Myc before and after zotiraciclib treatment. **G.** Promoters for *ACTB* and *GAPDH* show no Myc binding and are unaffected by zotiraciclib treatment. **H.** Model in which inhibition of CDK7 and CDK9 by zotiraciclib depletes Myc and Myc-driven transcriptional activity at both the *MYC* promoter and its cognate enhancers.

We then focused on the topologically associated domain (TAD) surrounding the *Myc* promoter, intersecting this data with H3K27ac ChIP-seq^35^, H3K4me1 CUT&RUN, and RNA-seq from D458 cells to refine active E-P contacts (**Figure 4C**, top). Of the 8 Myc-anchored loops conserved between both samples, 7 cognate anchors showed H3K4me1 deposition consistent with enhancer elements. Two of these lacked convincing H3K27ac or RNA-seq reads and thus may have been inactive in the cell line, while the remainder showed strong H3K27ac and active transcription. This included an enhancer locus ∼90kb upstream of the *Myc* promoter that showed exceptionally strong contact frequency and has been previously validated as a group 3 subgroup-specific enhancer^44^ (**Figure 4C**, *ENSG00000249375* left). When we examined Myc binding by CUT&RUN, we found each of the active putative enhancers to themselves be bound by Myc, consistent with the feed-forward mechanism commonly described in transcriptional core regulatory circuitry (CRC)^49^. Inspection of the patient-specific loop anchors showed variability in co-localized activating chromatin marks and RNA-seq reads, suggesting that many of these may also reflect functionally active if variable E-P loops (**Figure 4C**, *LINC00824* right).

Next, we tested whether zotiraciclib treatment could be used to disrupt transcriptional activity at these enhancers and, by extension, their collective amplification of transcription at the *Myc* promoter itself. We treated D458 cells with zotiraciclib (200 nM for 24 hours) before repeating the Myc CUT&RUN and RNA-seq experiments. At each of the loci identified with active enhancer features, treatment with zotiraciclib led to a loss of Myc occupancy and loss of transcriptional activity as measured by RNA-seq (**Figure 4C**, bottom). Consequently, at the *Myc* promotor we observed a dramatic loss of Myc binding and corresponding abrogation of transcriptional output (**Figure 4D**). This ZTR-induced loss of Myc expression was consistent when validated by qPCR and Western blot across multiple cell lines (**Figure 4E-F**). To assess whether this transcriptional depletion was a global phenomenon, we examined the same chromatin features at several housekeeping genes (*ACTB, GAPDH*). At baseline, these loci notably had neither E-P loops identified by Hi-C nor significant Myc binding by CUT&RUN, and when exposed to zotiraciclib, transcriptional output was comparatively unaffected (**Figure 4G**). In total, these data support a model in which CDK9/7i treatment potently inhibits E-P transcriptional activity within the *Myc*-amplified TAD, resulting in a marked loss of overall Myc expression (**Figure 4H**). The limitation in sample numbers subjected to Hi-C means that the characterization of E-P looping shown here is illustrative rather than comprehensive, and a much larger cohort would be required to robustly map the spectrum of enhancer structure for Myc-MB. However, the conserved role of CDK7 and CDK9 in modulating transcriptional output means that this therapeutic dependency could be reasonably anticipated to hold true irrespective of the specific structural arrangement within a given patient tumor.

### CDK9/7 Inhibition Suppresses Myc-driven Oncogenic Transcriptional Programs

Myc acts broadly as transcriptional amplifier, stimulating Pol II pause-release in both neoplastic and non-transformed cells^11,17^. Aberrant Myc activity drives oncogenic transcriptional programs involved in stemness, proliferation, and DNA repair across a range of cancers, including Myc-MB^34,50,51^. Given the depletion of Myc expression we observed following CDK9/7i treatment, we predicted a concordant suppression of downstream Myc-driven oncogenic functions. We first visualized this directly by combining immunofluorescence for Myc and nascent RNA synthesis. This demonstrated high levels of nuclear Myc localization uniformly across the Myc-MB cells, with puncta of nascent transcriptional activity clearly identifiable. Treatment with zotiraciclib led to a complete loss of appreciable Myc expression and significant downregulation of nascent transcription, though some foci of transcriptional activity persisted (**Figure 5A**). To characterize the consequences of this on Myc-driven transcriptional program, we examined changes to Myc binding (CUT&RUN, n=2) and gene expression (RNA-seq, n=3) genome-wide. This demonstrated a marked loss of Myc occupancy on chromatin following zotiraciclib treatment (**Figure 5B**). Differentially lost peaks were enriched for gene programs involved in chromatin remodeling and transcriptional regulation, cell cycle regulation, and DNA repair (**Figure 5C** and **Supplemental Table 3**). We replicated the RNA-seq experiment in an additional cell line (D425, n=3) and sorted for gene sets that were differentially enriched in both cell lines (n=3 replicates per model, p adj <0.05). Reproducibly enriched programs included downregulation of hallmark Myc signaling, cell cycle checkpoint regulation, and DNA repair, with a reciprocal upregulation of apoptosis (**Figure 5D-E** and **Supplemental Figure 4**). TRRUST analysis^52^ of transcriptional regulatory networks implicated in the observed expression changes clearly inferred Myc as the strongest candidate TF driving these programs (**Figure 5F** and **Supplemental Table 4**).

**Figure 5.**
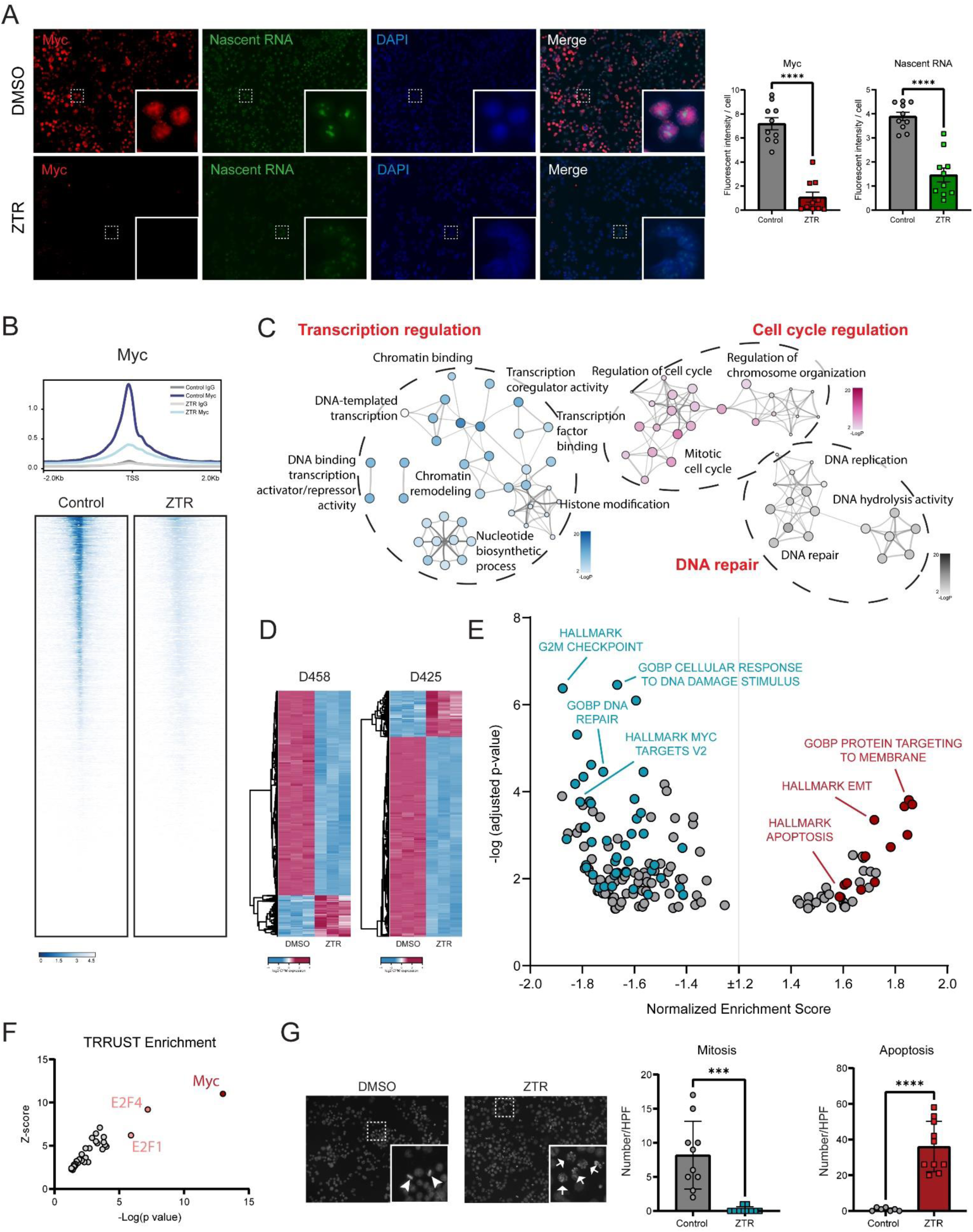
CDK9/7 inhibition suppresses Myc-driven oncogenic transcriptional programs. **A.** Click-IT fluorescent assay of relative nascent RNA abundance with immunofluorescent staining for Myc in the presence or absence of zotiraciclib treatment. Quantification of indicated fluorescent intensity on right, comparison reflects two-tailed Student’s t-test (**** p=<0.0001). **B.** Metagene profile (top) and heatmap (bottom) of Myc occupancy by CUT&RUN before and after zotiraciclib treatment (n=2 biological replicates). **C.** Gene ontology network constructed from genes with decreased Myc binding in (B). Each node denotes an enriched term, with color density reflecting -log(Pval). **D.** Unsupervised hierarchical clustering of D458 and D425 gene expression LFC ≥ 2 in control vs zotiraciclib-treated samples (n=3, p<0.001). **E.** Gene sets enriched (FDR padj <0.05) in (D). Colored circles (red=up, blue=down) denote gene sets significantly enriched in both cell models, while grey indicates terms unique to one model. **F.** TRRUST inference of transcription factor-target pairs from genes differentially downregulated following zotiraciclib treatment, ranked by Fisher’s exact test (-log(p val)). **G.** Greyscale of DAPI channel from (A) showing representative images (left) and quantification (right) of mitotic figures and apoptotic bodies in control vs zotiraciclib-treated samples. Comparison reflects two-tailed Student’s t-test (*** p=<0.001, **** p=<0.0001).

We then validated these transcriptomic results via functional assays. Quantification of direct microscopy demonstrated a significant decrease in mitotic figures with a corresponding increase in apoptotic bodies visible following zotiraciclib treatment (**Figure 5G**). Synchronized cell cycle analysis by flow cytometry showed a G1 arrest following ZTR exposure (**Supplemental Figure 5A**), with a significant induction of apoptosis as assessed by caspase 3 cleavage by immunoblot (**Supplemental Figure 5B**). Given the central role of ionizing radiation (IR) in standard-of-care multimodal therapy for Myc-MB^7^, we assessed IR-induced DNA double strand breaks (DSBs) as a surrogate for DNA damage response efficiency. ZTR treatment alone resulted in a significant accumulation in DSBs as measured by yH2AX, increased further when applied in combination with IR (**Supplemental Figure 6**). Collectively, these data establish that CDK9/7i-mediated Myc depletion leads to a loss of Myc binding on chromatin and a consequent loss of Myc-driven transcriptional programs. These programs represent core oncogenic functions in Myc-MB, including proliferation, DNA repair, and control of apoptosis. CDK9/7i therefor presents a novel, clinically applicable means of disrupting the core TF circuitry underpinning Myc-MB growth and survival.

### CDK9i Improves Survival in Flank but not Intracranial Xenograft Models of Myc-MB

As these data supported the utility of selective CDK9 inhibition as a therapeutic strategy for Myc-MB, we finally tested whether CDK9i would maintain anti-tumor effect *in vivo*. The agents utilized in this study have primarily been developed in the context of hematologic malignancies, and subsequent data has suggested that CNS exposure across the blood-brain barrier may be poor (^28,29^ and AstraZeneca, Investigator Brochure). Athymic nude mice were injected orthotopically into the cerebellum with D458 cells and randomized to control or AZD4573 treatment (15/15 mg/kg intraperitoneal biweekly^40^). Unfortunately, mice treated with AZD4573 showed no benefit in either survival or tumor volume as assessed by MRI (**Figure 6A**).

**Figure 6.**
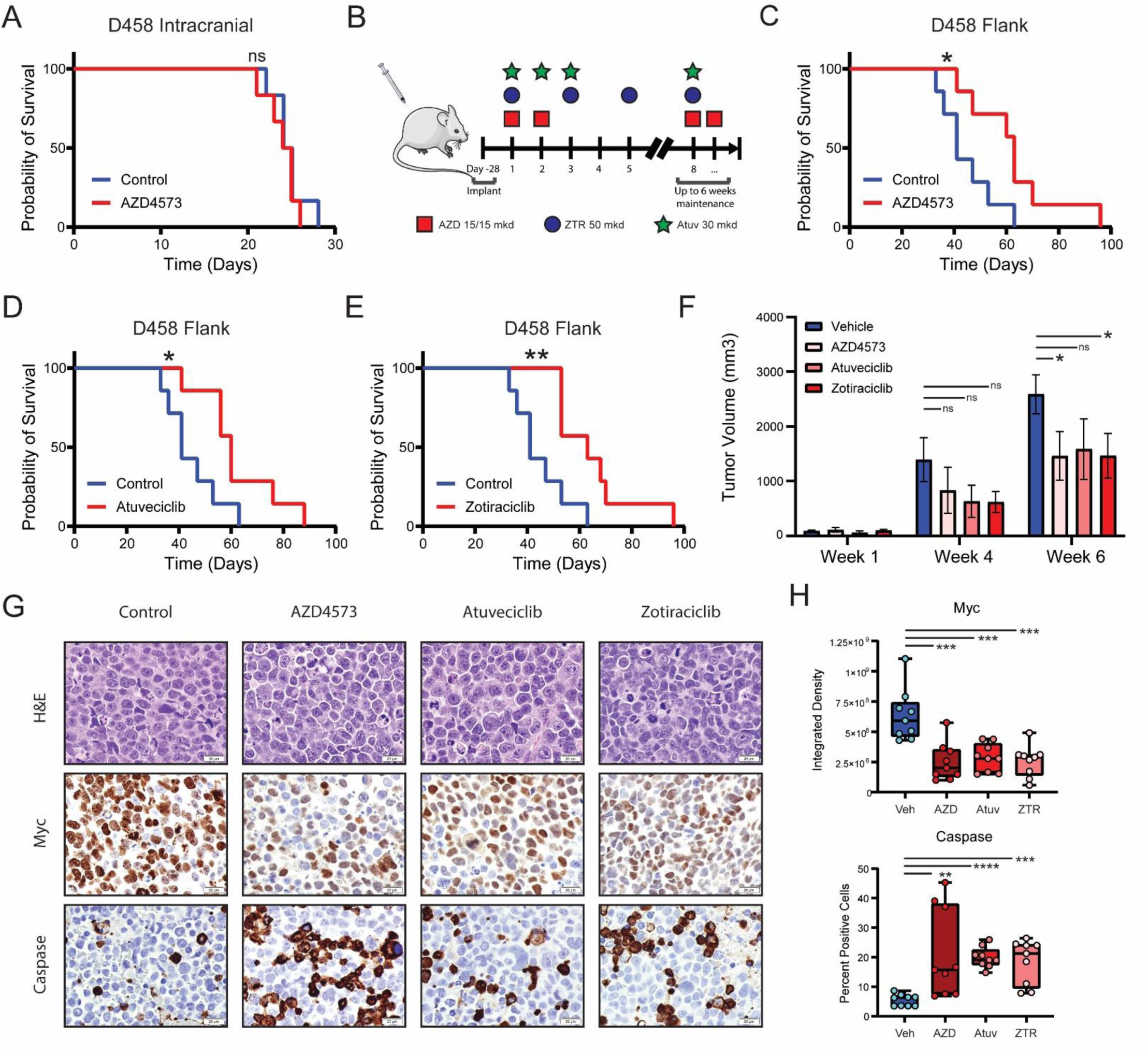
CDK9i improves survival in flank but not intracranial xenograft models of Myc-MB. **A.** Kaplan-Meier survival analysis of intracranial xenografts treated with AZD4573. **B.** Schema of treatment delivery for AZD4573, atuveciclib, or zotiraciclib in subcutaneous xenograft models. **C.** Kaplan-Meier survival analysis of subcutaneous xenografts treated with AZD4573 at same dosing as (A). Kaplan-Meier survival analysis of subcutaneous xenografts treated with atuveciclib (**D**) or zotiraciclib (**E**). **F.** Tumor volumes from (C-E). Comparisons reflect pairwise two-tailed Student’s t-tests (* p=<0.05). **G.** Representative immunohistochemistry for Myc and cleaved caspase expression in tumors harvested from (C-E). Scale bar represents 20 µm. **H.** Quantification of immunohistochemistry from (G). Comparison reflects two-tailed Student’s t-test (** p=<0.01, *** p=<0.001, **** p=<0.0001), n=3 animals per arm with n=3 high-powered fields per animal.

To assess anti-tumor effect of CDK9 inhibition *in vivo* independent of the CNS penetration of a specific agent, we broadened testing of CDK9i using subcutaneous models. Animals were injected in the flank and randomized to either AZD4573 (15/15 mg/kg intraperitoneal biweekly^40^), atuveciclib (30 mg/kg/dose daily oral gavage; 3 days on / 2 days off^28^), or zotiraciclib (50 mg/kg oral gavage three times weekly)(**Figure 6B**). In each cohort, treatment with CDK9i led to a statistically significant improvement in xenograft survival (median survival 41 vs 63 days AZD4573, p=0.03; vs 60 days atuveciclib, p=0.03; vs 63 days zotiraciclib, p=0.008) (**Figure 6C-E**). Tumor volumes were decreased in CDK9i-treated animals, with a concordant decrease in Myc intensity by IHC (**Figure 6F-G**). Our data here is consistent with prior reports of limited CNS exposures for this class of agents, and it highlights that while CDK9 is a relevant pharmacologic target for Myc-MB, consideration of pharmacokinetic properties will remain important for effective application to primary CNS disease.

## Discussion

Myc has been studied has a master oncogenic transcription factor for decades^50,51^. In most core regulatory circuitry, binding of TFs to their own enhancers amplifies transcription of the TF itself and subsequently the downstream transcriptional networks under that TF control, enforcing core programs of cell identity^49^. In states of neoplastic transformation, this regulatory circuitry can be co-opted to drive oncogenic functions. Tightly controlled, stepwise enhancer activation and chromatin looping similarly allows for ordered transcriptional patterning across development, but it too can be highly disordered in cancer, even more so when structural or copy number variants rearrange regulatory regions directly^48,53^. Our Hi-C data here highlights this variability, with considerable diversity of E-P looping even to a single promoter when compared between patients. As a result, targeting the transcriptional cofactors required at each loci agnostic of higher-order structure may represent a more reproducible therapeutic strategy.

Small molecule inhibitors targeting transcriptional cofactors have shown promise as therapeutic means of drugging enhancer-dependent oncogenes^37,38^. While many of the early clinical trials have been conducted primarily in hematologic malignancies^39,40^, our group and others have highlighted these cofactors as relevant targets in CNS disease^28,29,34,54^. In this present study, we leverage this understanding of Myc CRC to identify and apply compounds targeting the transcriptional cofactors CDK9 and CDK7 as a treatment strategy for Myc-MB. CDK9 and CDK7 are conserved dependencies across models of Myc-MB, and inhibiting these kinases leads to a marked depletion of Myc expression. This in turn causes Myc-driven oncogenic programs to collapse, exerting a significant anti-tumor effect both *in vitro* and *in vivo*. Ongoing development of CDK9 inhibitors would benefit from pharmacologic consideration of CNS exposure in order to realize the potential for Myc-directed therapies in high-risk CNS disease.

## Author Contributions

L.M.S. and D.W. performed the CUT&RUN and RNA-seq experiments. L.M.S. and Arima Genomics performed the Hi-C experiments. E.D. and Pluto performed bioinformatic analysis. K.M., F.M.W., and S.D. performed in vitro phenotypic assays. A.P. and F.M.W performed the stereotactic xenograft injections; F.M.W., K.M., and I.B. conducted the subsequent in vivo studies. M.M.H and B.J.C performed patient FISH and amplification analysis. N.J.S. performed the xenograft MR imaging and was responsible for its analysis. N.K.F. procured tissue specimens and maintained IRB compliance. F.M.W., K.M., L.M.S., and N.A.D. prepared the figures and wrote the manuscript. S.V., B.V., R.V., and N.A.D. conceived the project, supervised all aspects of the work, and edited the manuscript.

## Acknowledgements

D425 and D458 cell lines were generously provided by Dr. Darrel D. Bigner (Duke University Medical Center, NC). The ONS-76 medulloblastoma cell line was kindly given by Dr. James T. Rutka (University of Toronto, Canada) and the UW228 cell line by Dr. John Silber (University of Washington, Seattle). We likewise thank the University of Colorado Functional Genomics Facility, Genomics Shared Resource, Research Histology Shared Resource, Bioinformatics and Biostatistics Shared Resource, and Animal Imaging Shared Resource for their contributions to this work. Original scientific illustrations by Katie Vicari.

## Data Availability

The accession number for the raw and processed data reported in this paper is GEO: GSE267473.

