## Supplementary Data for "Targeting processive transcription for Myc-driven circuitry in medulloblastoma"

**Supplementary Table 1.** CRISPR-Cas9 screens in Myc-MB.

**Supplementary Table 2.** Hi-C loops anchored at Myc promoter.

**Supplementary Table 3.** Myc CUT&RUN enrichment following zotiraciclib treatment.

**Supplementary Table 4.** TRRUST enrichment following zotiraciclib treatment.

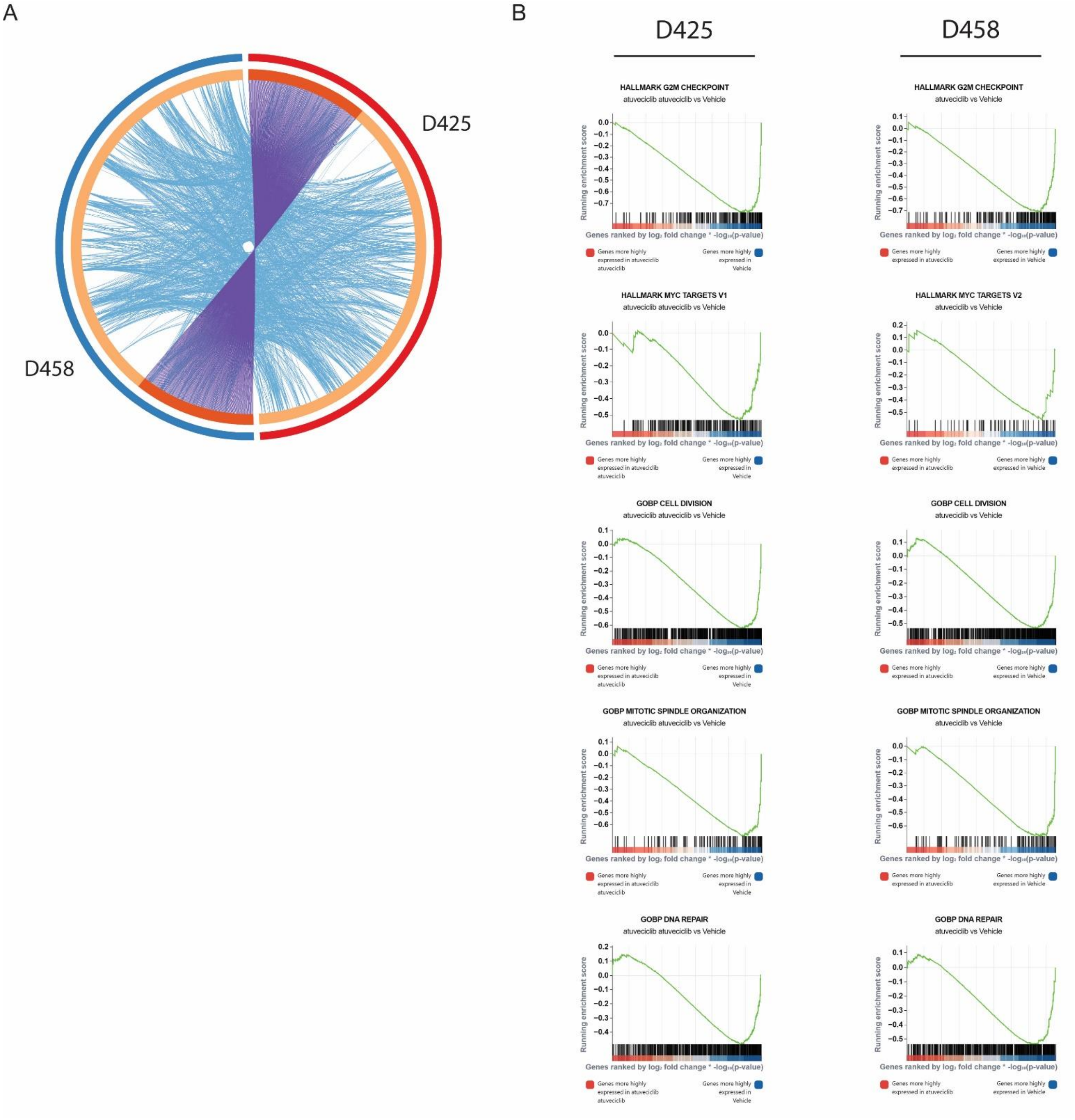


**Supplemental Figure 1.** Atuveciclib-induced transcriptional changes display strong overlap between models. **A.** Circlos plot of genes differentially upregulated (top 1000 genes, p adj <0.05) in D425 and D458 models. Purple connections indicate identical genes in both lists, while blue connections indicate common enriched GO terms. **B.** Representative common gene sets from RNA-seq in D425 (left) and D458 (right) models.

A

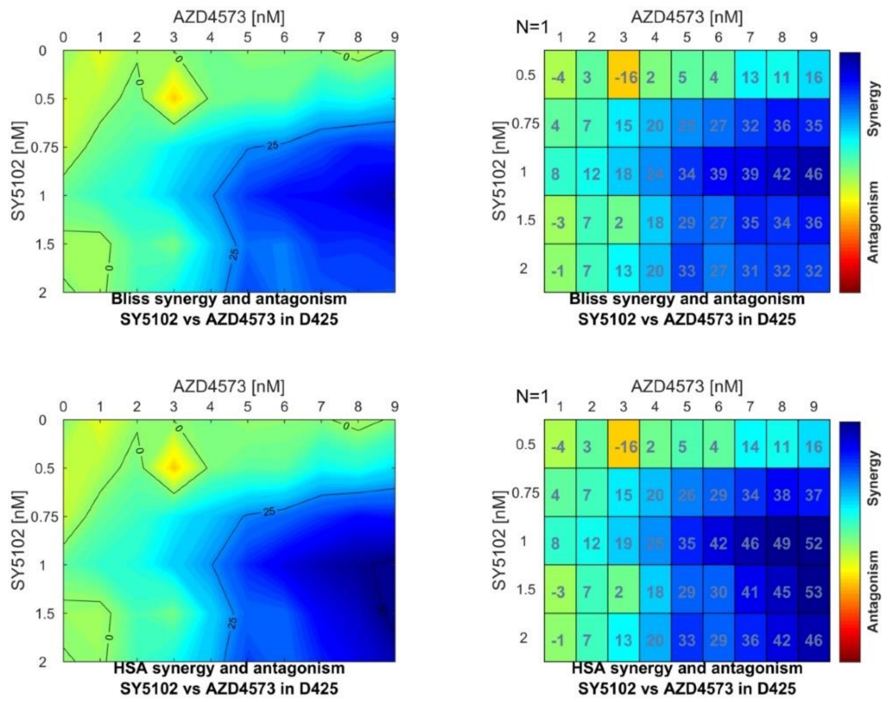


B

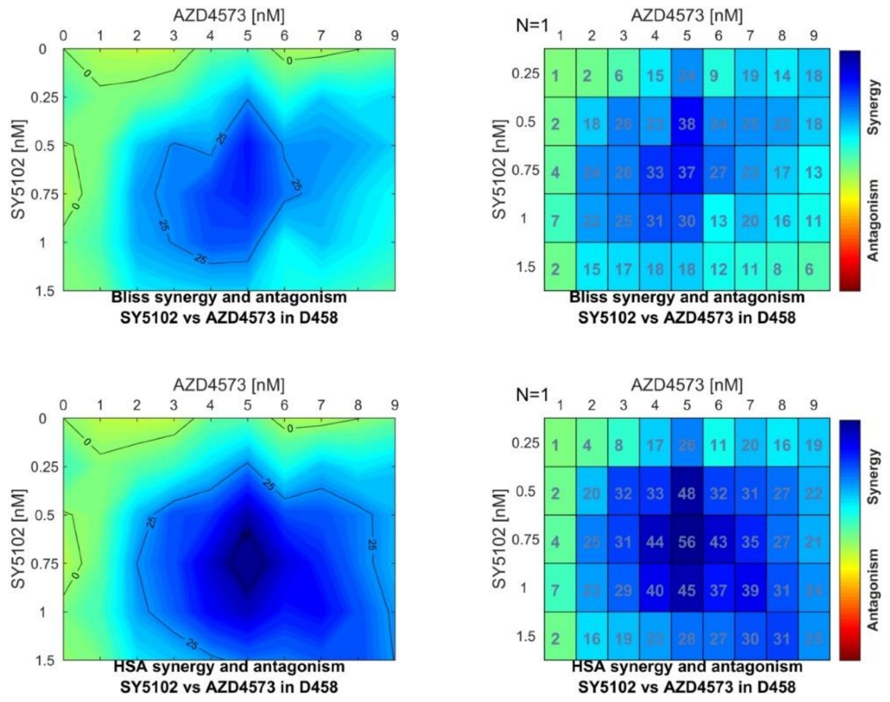


**Supplemental Figure 2.** Selective CDK9i and CDK7i are highly synergistic. **A.** Bliss model (top) and HSA model (bottom) of synergy scoring at increasing concentrations of AZD4573 and SY5102 in D425 cells. Scores ≥10 are considered synergistic, represented as cyan to blue. **B.** Bliss model (top) and HSA model (bottom) of synergy scoring at increasing concentrations of AZD4573 and SY5102 in D458 cells. Scores ≥10 are considered synergistic, represented as cyan to blue.

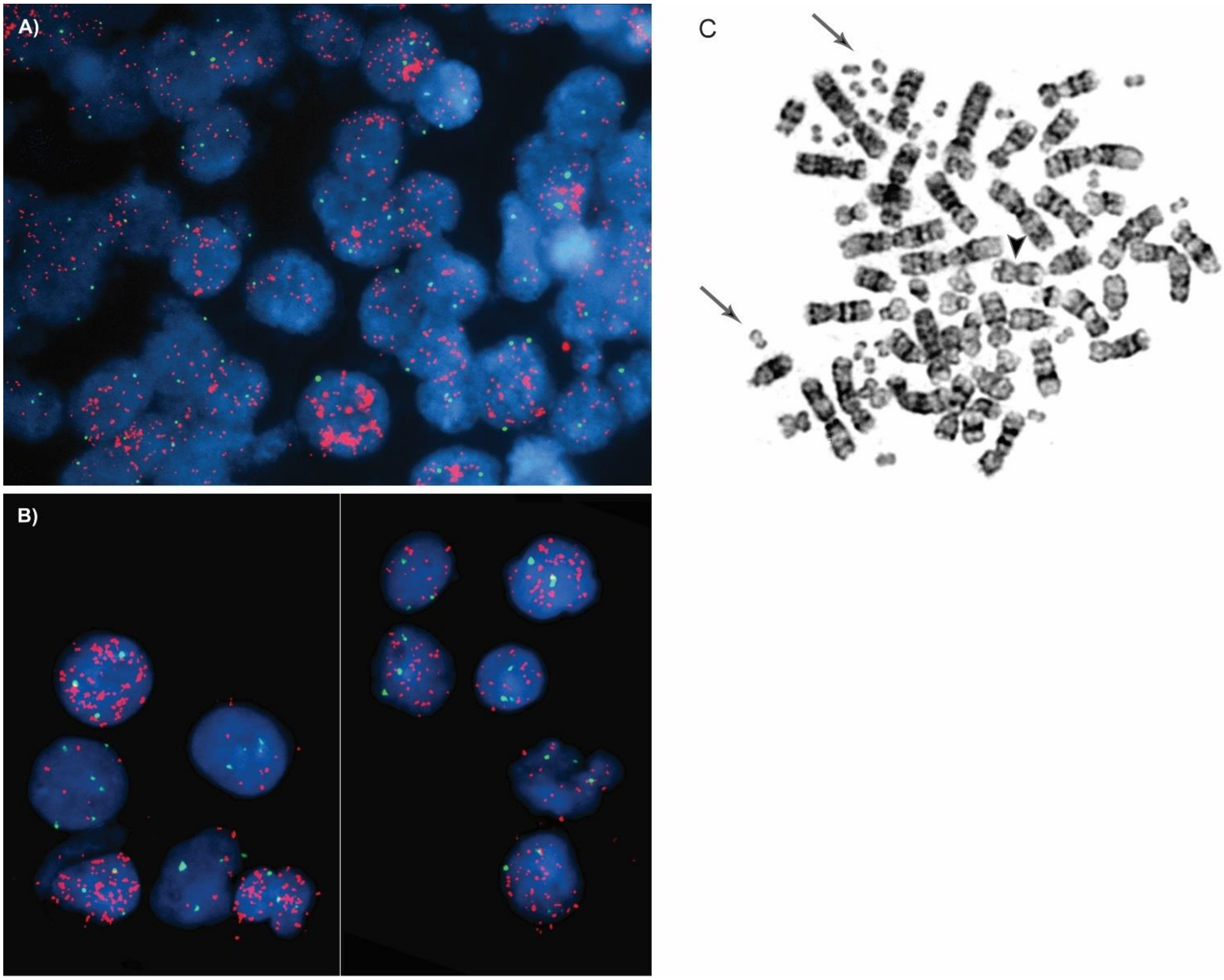


**Supplemental Figure 3.** Clinical annotation of Myc amplification in primary patient samples. **A.** and **B.** Interphase FISH analysis on brain tissue was performed using probes for chromosome 8 centromere (green signal) and MYC at locus 8q24 (red signal) (Vysis, Abbott Molecular). Top and bottom panels both display a high level MYC amplification in the majority of cells in a pattern consistent with double minutes. **C.** Metaphase of G-banded chromosomes from patient in (B) demonstrating multiple double minutes (arrows) and an isochromosome 17q (arrowhead).

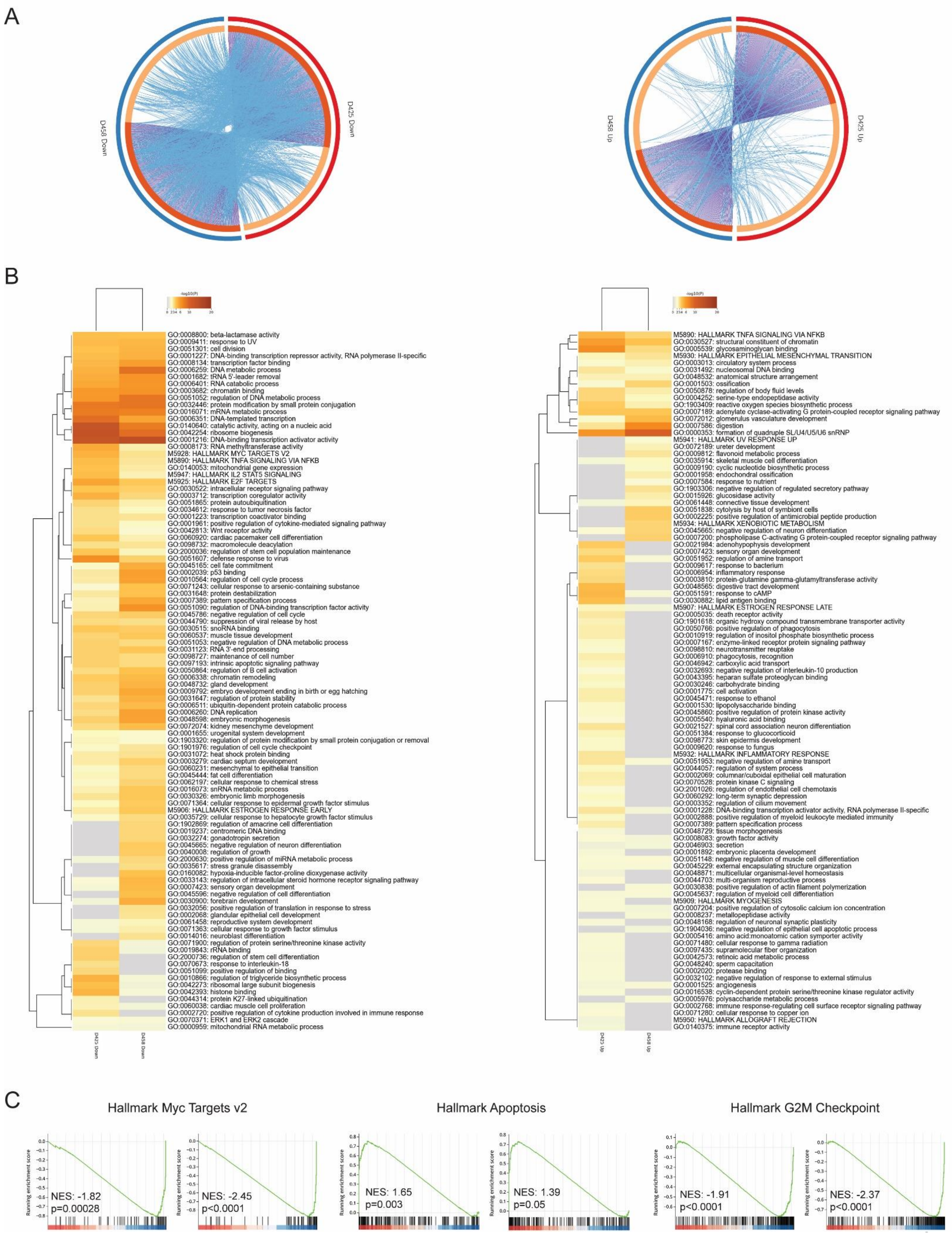
A
D458 Down
D425 Down
D458 Up
D425 Up
B
-log10(P)
0 2 3 4 6 10 20
GO:0008800: beta-lactamase activity
GO:0009411: response to UV
GO:0051301: cell division
GO:0001227: DNA-binding transcription repressor activity, RNA polymerase II-specific
GO:0008134: transcription factor binding
GO:0006259: DNA metabolic process
GO:0001682: tRNA 5'-leader removal
GO:0006401: RNA catabolic process
GO:0003682: chromatin binding
GO:0051052: regulation of DNA metabolic process
GO:0032446: protein modification by small protein conjugation
GO:0016071: mRNA metabolic process
GO:0006351: DNA-templated transcription
GO:0140640: catalytic activity, acting on a nucleic acid
GO:0042254: ribosome biogenesis
GO:0001216: DNA-binding transcription activator activity
GO:0008173: RNA methyltransferase activity
M5928: HALLMARK MYC TARGETS V2
M5890: HALLMARK TNFA SIGNALING VIA NFKB
GO:0140053: mitochondrial gene expression
M5947: HALLMARK IL2 STAT5 SIGNALING
M5925: HALLMARK E2F TARGETS
GO:0030522: intracellular receptor signaling pathway
GO:0003712: transcription coregulator activity
GO:0051865: protein autoubiquitination
GO:0034612: response to tumor necrosis factor
GO:0001223: transcription coactivator binding
GO:0001961: positive regulation of cytokine-mediated signaling pathway
GO:0042813: Wnt receptor activity
GO:0060920: cardiac pacemaker cell differentiation
GO:0098732: macromolecule deacylation
GO:2000036: regulation of stem cell population maintenance
GO:0051607: defense response to virus
GO:0045165: cell fate commitment
GO:0002039: p53 binding
GO:0010564: regulation of cell cycle process
GO:0071243: cellular response to arsenic-containing substance
GO:0031648: protein destabilization
GO:0007389: pattern specification process
GO:0051090: regulation of DNA-binding transcription factor activity
GO:0045786: negative regulation of cell cycle
GO:0044790: suppression of viral release by host
GO:0030515: snoRNA binding
GO:0060537: muscle tissue development
GO:0051053: negative regulation of DNA metabolic process
GO:0031123: RNA 3'-end processing
GO:0098727: maintenance of cell number
GO:0097193: intrinsic apoptotic signaling pathway
GO:0050864: regulation of B cell activation
GO:0006338: chromatin remodeling
GO:0048732: gland development
GO:0009792: embryo development ending in birth or egg hatching
GO:0031647: regulation of protein stability
GO:0006511: ubiquitin-dependent protein catabolic process
GO:0006260: DNA replication
GO:0048598: embryonic morphogenesis
GO:0072074: kidney mesenchyme development
GO:0001655: urogenital system development
GO:1903320: regulation of protein modification by small protein conjugation or removal
GO:1901976: regulation of cell cycle checkpoint
GO:0031072: heat shock protein binding
GO:0003279: cardiac septum development
GO:0060231: mesenchymal to epithelial transition
GO:0045444: fat cell differentiation
GO:0062197: cellular response to chemical stress
GO:0016073: snRNA metabolic process
GO:0030326: embryonic limb morphogenesis
GO:0071364: cellular response to epidermal growth factor stimulus
M5906: HALLMARK ESTROGEN RESPONSE EARLY
GO:0035729: cellular response to hepatocyte growth factor stimulus
GO:1902869: regulation of amacrine cell differentiation
GO:0019237: centromeric DNA binding
GO:0032274: gonadotropin secretion
GO:0045665: negative regulation of neuron differentiation
GO:0040008: regulation of growth
GO:2000630: positive regulation of miRNA metabolic process
GO:0035617: stress granule disassembly
GO:0160082: hypoxia-inducible factor-proline dioxygenase activity
GO:0033143: regulation of intracellular steroid hormone receptor signaling pathway
GO:0007423: sensory organ development
GO:0045596: negative regulation of cell differentiation
GO:0030900: forebrain development
GO:0032056: positive regulation of translation in response to stress
GO:0002068: glandular epithelial cell development
GO:0061458: reproductive system development
GO:0071363: cellular response to growth factor stimulus
GO:0014016: neuroblast differentiation
GO:0071900: regulation of protein serine/threonine kinase activity
GO:0019843: rRNA binding
GO:2000736: regulation of stem cell differentiation
GO:0070673: response to interleukin-18
GO:0051099: positive regulation of binding
GO:0010866: regulation of triglyceride biosynthetic process
GO:0042273: ribosomal large subunit biogenesis
GO:0042393: histone binding
GO:0044314: protein K27-linked ubiquitination
GO:0060038: cardiac muscle cell proliferation
GO:0002720: positive regulation of cytokine production involved in immune response
GO:0070371: ERK1 and ERK2 cascade
GO:0000959: mitochondrial RNA metabolic process
D425 Down
D458 Down
M5890: HALLMARK TNFA SIGNALING VIA NFKB
GO:0030527: structural constituent of chromatin
GO:0005539: glycosaminoglycan binding
M5930: HALLMARK EPITHELIAL MESENCHYMAL TRANSITION
GO:0003013: circulatory system process
GO:0031492: nucleosomal DNA binding
GO:0048532: anatomical structure arrangement
GO:0001503: ossification
GO:0050878: regulation of body fluid levels
GO:0004252: serine-type endopeptidase activity
GO:1903409: reactive oxygen species biosynthetic process
GO:0007189: adenylate cyclase-activating G protein-coupled receptor signaling pathway
GO:0072012: glomerulus vasculature development
GO:0007586: digestion
GO:0000353: formation of quadruple SL/U4/U5/U6 snRNP
M5941: HALLMARK UV RESPONSE UP
GO:0072189: ureter development
GO:0009812: flavonoid metabolic process
GO:0035914: skeletal muscle cell differentiation
GO:0009190: cyclic nucleotide biosynthetic process
GO:0001958: endochondral ossification
GO:0007584: response to nutrient
GO:1903306: negative regulation of regulated secretory pathway
GO:0015926: glucosidase activity
GO:0061448: connective tissue development
GO:0051838: cytolysis by host of symbiont cells
GO:0002225: positive regulation of antimicrobial peptide production
M5934: HALLMARK XENOBIOTIC METABOLISM
GO:0045665: negative regulation of neuron differentiation
GO:0007200: phospholipase C-activating G protein-coupled receptor signaling pathway
GO:0021984: adenohypophysis development
GO:0007423: sensory organ development
GO:0051952: regulation of amine transport
GO:0009617: response to bacterium
GO:0006954: inflammatory response
GO:0003810: protein-glutamine gamma-glutamyltransferase activity
GO:0048565: digestive tract development
GO:0051591: response to cAMP
GO:0030882: lipid antigen binding
M5907: HALLMARK ESTROGEN RESPONSE LATE
GO:0005035: death receptor activity
GO:1901618: organic hydroxy compound transmembrane transporter activity
GO:0050766: positive regulation of phagocytosis
GO:0010919: regulation of inositol phosphate biosynthetic process
GO:0007167: enzyme-linked receptor protein signaling pathway
GO:0098810: neurotransmitter reuptake
GO:0006910: phagocytosis, recognition
GO:0046942: carboxylic acid transport
GO:0032693: negative regulation of interleukin-10 production
GO:0043395: heparan sulfate proteoglycan binding
GO:0030246: carbohydrate binding
GO:0001775: cell activation
GO:0045471: response to ethanol
GO:0001530: lipopolysaccharide binding
GO:0045860: positive regulation of protein kinase activity
GO:0005540: hyaluronic acid binding
GO:0021527: spinal cord association neuron differentiation
GO:0051384: response to glucocorticoid
GO:0098773: skin epidermis development
GO:0009620: response to fungus
M5932: HALLMARK INFLAMMATORY RESPONSE
GO:0051953: negative regulation of amine transport
GO:0044057: regulation of system process
GO:0002069: columnar/cuboidal epithelial cell maturation
GO:0070528: protein kinase C signaling
GO:2001026: regulation of endothelial cell chemotaxis
GO:0060292: long-term synaptic depression
GO:0003352: regulation of cilium movement
GO:0001228: DNA-binding transcription activator activity, RNA polymerase II-specific
GO:0002888: positive regulation of myeloid leukocyte mediated immunity
GO:0007389: pattern specification process
GO:0048729: tissue morphogenesis
GO:0008083: growth factor activity
GO:0046903: secretion
GO:0001892: embryonic placenta development
GO:0051148: negative regulation of muscle cell differentiation
GO:0045229: external encapsulating structure organization
GO:0048871: multicellular organismal-level homeostasis
GO:0044703: multi-organism reproductive process
GO:0030838: positive regulation of actin filament polymerization
GO:0045637: regulation of myeloid cell differentiation
M5909: HALLMARK MYOGENESIS
GO:0007204: positive regulation of cytosolic calcium ion concentration
GO:0008237: metallopeptidase activity
GO:0048168: regulation of neuronal synaptic plasticity
GO:1904036: negative regulation of epithelial cell apoptotic process
GO:0005416: amino acid:monoatomic cation symporter activity
GO:0071480: cellular response to gamma radiation
GO:0097435: supramolecular fiber organization
GO:0042573: retinoic acid metabolic process
GO:0048240: sperm capacitation
GO:0002020: protease binding
GO:0032102: negative regulation of response to external stimulus
GO:0001525: angiogenesis
GO:0016538: cyclin-dependent protein serine/threonine kinase regulator activity
GO:0005976: polysaccharide metabolic process
GO:0002768: immune response-regulating cell surface receptor signaling pathway
GO:0071280: cellular response to copper ion
M5950: HALLMARK ALLOGRAFT REJECTION
GO:0140375: immune receptor activity
D425 Up
D458 Up
C
Hallmark Myc Targets v2
Hallmark Apoptosis
Hallmark G2M Checkpoint
Running enrichment score
NES: -1.82
p=0.00028
NES: -2.45
p<0.0001
NES: 1.65
p=0.003
NES: 1.39
p=0.05
NES: -1.91
p<0.0001
NES: -2.37
p<0.0001

**Supplemental Figure 4.** Zotiraciclib-induced transcriptional changes display strong overlap between models. **A.** Circlos plot of differentially down- (left) or upregulated (right) genes (top 1000 genes, p adj <0.05) in D425 and D458 models. Purple connections indicate identical genes in both lists, while blue connections indicate common enriched GO terms. **B.** Clustered heatmap of GO terms enriched from (A). **C.** Representative common gene sets from RNA-seq in D425 (left) and D458 (right) models.

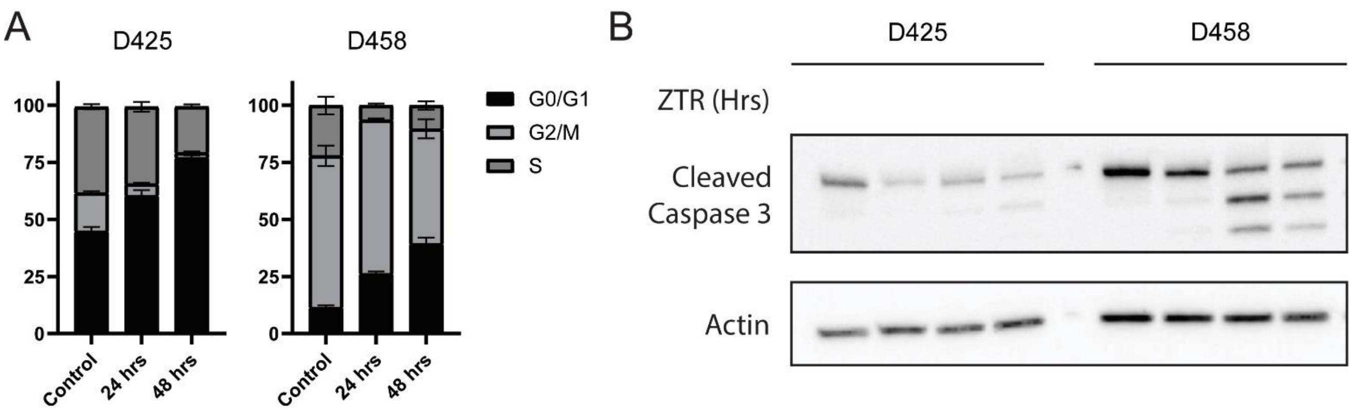


**Supplemental Figure 5.** Validation of cell cycle and apoptosis. **A.** Cell cycle analysis of D425 and D458 cells over time in the presence of zotiraciclib. **B.** Immunoblot for cleaved caspase 3 at indicated timepoints following zotiraciclib treatment.

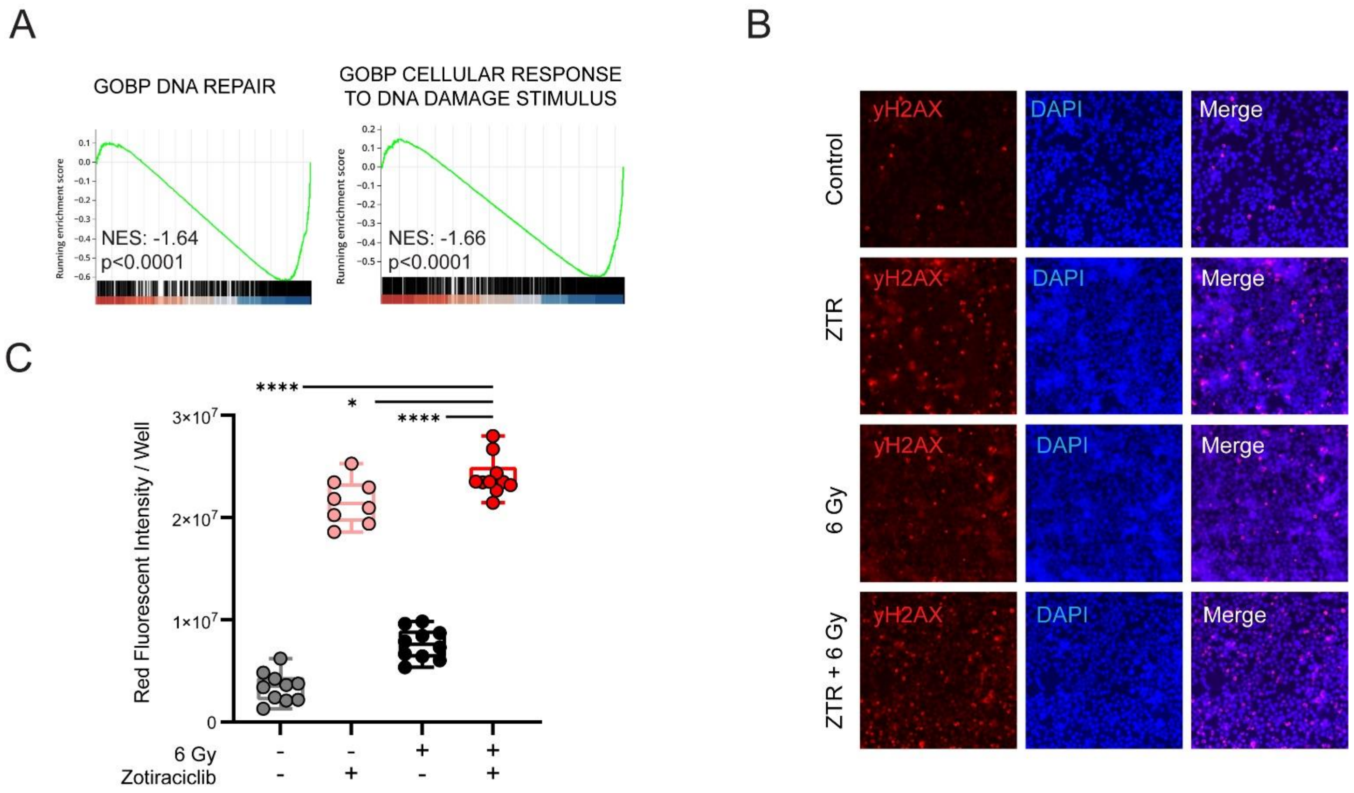


**Supplemental Figure 6.** Zotiraciclib treatment impairs DNA damage response programs. **A.** Gene sets related to DNA repair following zotiraciclib treatment. **B.** Immunofluorescent staining for yH2AX in D458 cells following zotiraciclib treatment, 6 Gy radiation, or combination. **C.** Quantification from imaging in (B).
